# Breast cancer cells can recognize and respond to different levels of progestins to achieve different phenotypic outputs

**DOI:** 10.1101/2023.10.13.562206

**Authors:** Emma L. Dolan, Rachid Safi, Chingyi Chang, Hilary E. Eaton, Shweta Krishnan, Roya Safi, Paige Watkinson, Henry Long, Yingtian Xie, Myles Brown, Susan K. Murphy, Donald P. McDonnell

## Abstract

The steroid hormone progesterone, acting through its nuclear progesterone receptor (PR), has complex physiologic activities with different levels of hormones manifesting distinct and sometimes opposing phenotypic responses in target tissues. However, most of what is currently known about the transcriptional activity of PR comes from studies performed using progestins at levels that are in the high physiologic range (≥10nM), relevant only in the luteal phase of the reproductive cycle and pregnancy in humans. These studies do not consider the non-linearity of responses to progestins that exist in physiology and are not informative as to the mechanisms by which low levels of progestins, as occurs during menopause, exert their biological activities. Thus, we undertook to define the mechanisms which enable cells to recognize and respond to different levels of progestins. Using a PR expressing cell model of luminal breast cancer (T47D cells) we demonstrated that low concentration progestins (0.1-0.3nM) drive proliferation while high dose progestins (≥10nM) inhibit proliferation. Using both unbiased and targeted approaches, we found that low dose progestins facilitate cell cycle entry by enhanced expression of CCND1 and SGK1, which are both required to initiate a signaling cascade that leads to increased phospho-Rb and E2F1 transcriptional activity. CCND1 cooperates with CDK4/6 to phosphorylate Rb, while SGK1 phosphorylates p21, thereby excluding it from the nucleus and inhibiting its anti-proliferative function. Expression of *CCND1* and *SGK1* mRNAs are primary responses to low dose progestin treatment. However, these responses occur at very low levels of receptor occupancy and in the absence of receptor phosphorylation events that have been shown to be required for nuclear translocation and transcriptional activity. These findings challenge the assumption of linearity in response to progestin dose. Further, they suggest that concentrations of progestins found in post-menopausal women (0.1-0.3nM) have the potential to exert proliferative responses in PR expressing cancers.

## INTRODUCTION

The steroid hormone progesterone (P4) is involved in the establishment and maintenance of reproductive function, having key roles in promoting breast development, regulating uterine cycling, supporting placentation during pregnancy, stimulating lactation, and suppressing immune function (1-4). The biological actions of progestins are restricted to cells that express specific high a?nity progesterone receptors (PR). In humans, differential promoter usage within the *PGR* gene results in the generation of two transcripts that encode functionally distinct receptor isoforms PR-A and PR-B. The resulting proteins are structurally identical except that the larger PR-B isoform includes an additional 164 amino acids at its amino terminus. While each receptor exhibits the same a?nity for ligands, they have been shown to regulate distinct transcriptomes and to have different functions in reproduction (5-7). A key function of PR-A is to negatively regulate PR-B activity and to dampen the transcriptional activity of the estrogen receptor (ER) (7, 8). In the absence of an activating ligand, both PR isoforms reside in the cytoplasm associated with a large heat shock protein (HSP) complex which maintains the receptor(s) in a transcriptionally inactive state (9). Upon ligand binding, the receptor(s) undergo an activating conformational change that results in its ejection from the HSP complex (10). Subsequently, PR-A and PR-B can form either homo or heterodimers which translocate to the nucleus, associate with specific DNA response elements within the enhancers of target genes and exert positive or negative effects on gene transcription (10). The transcriptional output of the DNA-bound receptor dimer is influenced by its ability to engage functionally distinct coregulators in target cells and is influenced by other signaling pathways that converge upon the receptor and receptor-associated coregulator complexes (11). It is also well established that differences in the response of different cells to progestins can be attributed to variations in the relative expression of PR-A and PR-B and to differences in the expression of functionally distinct coregulators (6).

A significant unanswered question in the field of PR pharmacology Is how differences in the levels of progestins, as occurs through various stages of the reproductive cycle in females, result in different functional outputs. For instance, during development and following menarche, reproductive organs recognize and respond to different levels of progestins to generate specific phenotypic changes that do not follow a classic linear dose-response relationship. Indeed, low levels of circulating P4 (<3nM), such as occurs in the follicular phase of the uterine cycle, results in increased endometrial proliferation and vascularization, while high levels of P4 (30-100nM) in the luteal phase of the cycle facilitates endometrial thickening and increased secretory activity (2). Additionally, during early pregnancy when hormone levels are low, P4 supports embryonic cell proliferation and breast epithelial expansion (12, 13). As pregnancy progresses and P4 levels rise (>300nM), its proliferative activities are not apparent but rather it promotes fetal cell differentiation, myometrial quiescence, and breast alveolar cell differentiation (13-16). These and other examples of the bifunctional nature of progestins suggest that some cells possess the ability to detect and respond in a distinct manner to different levels of this hormone.

PR expression has long been used as a prognostic measure of response to ER-directed endocrine therapy in breast cancer (17). However, even before the emergence of contemporary ER-directed endocrine therapies, high dose progestins, notably medroxyprogesterone acetate (MPA), were used as second/third line endocrine therapies in patients with advanced breast cancer (18). Other than a few unsuccessful attempts to develop antiprogestins for breast and gynecological cancers, PR remains an underexplored therapeutic target (19, 20). Indeed, it remains to be determined if PR-directed drugs should target the low dose or high dose biology of progestins, eliminate PR signaling completely, or whether there would be utility in developing drugs that exhibit process selectivity. The idea that absolute inhibition of PR signaling is a desired outcome is supported by large epidemiologic studies of menopausal hormone replacement therapy (HRT) which has indicated that progestins are mitogenic in the post-menopausal breast (21-27). However, this is at odds with what is known about the normal physiological function of progestins to oppose estrogen action in the reproductive tract and does not consider the fact that the response to this hormone changes as a function of dose.

It is clear that the pharmaceutical exploitation of PR has been limited by our lack of understanding of the mechanisms by which PR biology is impacted by the levels of progestins and by the assumption that absolute inhibition of progestin signaling should be the goal of PR-directed dugs for breast and gynecological cancers. This puts into context the results presented herein which suggest that low dose, but not high dose, progestins support cell proliferation.

## RESULTS

### Progestins drive breast cancer cell proliferation in a non-linear dose-responsive manner

It is generally assumed that cellular responses to progestins occur in a dose proportional manner with different levels of the hormone eliciting quantitative rather than qualitative changes in gene expression. However, the substantially different phenotypic responses induced by different levels of this hormone in the reproductive system, for instance, led us to explore the alternative hypothesis that response to progestins can occur in a non-linear manner. To explore this possibility, we evaluated the impact of different levels of progestins on breast cancer cell proliferation *in vitro*. In most ER+ breast cancer cell lines, estrogen signaling is required to induce and maintain PR expression (28). However, for this study we used the well validated human breast carcinoma cell line T47D as a model system, as PR expression is constitutive and uncoupled from estrogen signaling (29). Synchronized T47D cells were plated in charcoal stripped (CFS) media to achieve a condition of extreme hormone deprivation. Cells were then treated with increasing doses (0.03-100nM) of the synthetic progestin promegestone (R5020), and growth responses were recorded after 7 days. In this manner, it was determined that proliferation was induced by low doses (0.03-0.3nM) of R5020, while higher levels of the progestin were without effect (**Figure 1A**). Similar non-linear dose responses were observed upon treatment of cells with natural progesterone (P4) and with synthetic medroxyprogesterone acetate (MPA) (a clinically used synthetic progestin). Whereas high dose progestins did not induce proliferation in CFS, we observed that they were able to quantitatively inhibit the proliferation of cells grown in growth factor- and hormone-containing fetal bovine serum (FBS) media (**Figure 1B**). Further, 17β-estradiol (E2)-induced proliferation was completely inhibited by high dose progestins (**Figure 1C**). This finding is notable given the established clinical utility of high dose MPA as a treatment for metastatic ER+ breast cancer (18).

**Figure 1.**
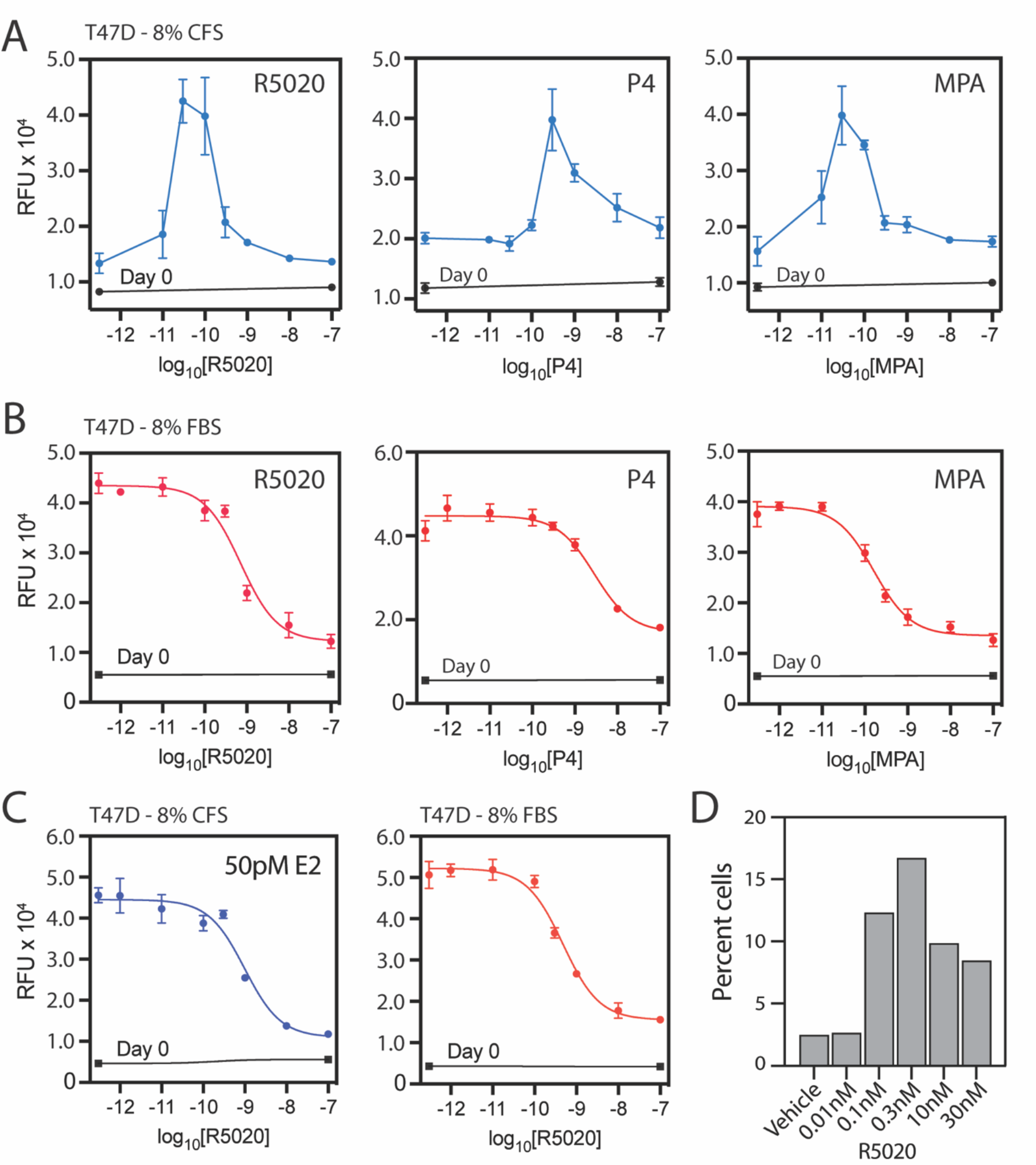
Low dose progestin treatment increases T47D breast cancer proliferation. Synchronized T47D cells, grown in (A) charcoal stripped media (CFS) or (B) fetal bovine serum (FBS), were treated with promegestone (R5020), natural progesterone (P4), or medroxyprogesterone acetate (MPA). Following 7 days of treatment, cell number was quantified by Hoechst DNA stain. Results are expressed as mean relative fluorescent units ± standard deviation. (C) High dose progestin treatment opposes 17β-estradiol-mediated proliferation. Synchronized T47D cells, grown in CFS in the presence of 50pM 17b-estradiol (E2) or FBS (for comparative purposes), were treated with different doses of R5020. Following 7 days of treatment, cell number was quantified by Hoechst DNA stain. Results are expressed as mean relative fluorescent units ± standard deviation. (D) Low dose progestins drive cell cycle entry. Synchronized T47D cells, grown in CFS, were treated with different doses of R5020 for 18hr. Cells were fixed and stained with DNA stain propidium iodide and assessed by flow cytometry. Results are expressed as percent cell population in S-phase of the cell cycle.

Cell cycle analysis revealed that the effects of different doses of progestins on proliferation represented changes in the number of cells in S-phase, as opposed to cell death. Specifically, synchronized T47D cells were treated with a range of doses of R5020 for 18hr and the number of cells in S-phase were enumerated by flow cytometry. In this manner it was determined that low dose R5020 (0.1-0.3nM) increased entry into S-phase, an activity which was considerably reduced upon treatment with higher doses of R5020 (**Figure 1D**). These findings are consistent with other studies which demonstrated that high dose progestins can induce transient cell cycle progression prior to facilitating cell cycle arrest in G1 (30, 31). Taken together, these studies have uncovered an unexpected complexity in PR pharmacology revealing that cells can distinguish between different levels of hormone to accomplish different phenotypic outputs.

### Low dose progestins induce a unique transcriptional landscape enriched with cell cycle regulators

To identify the genes responsible for the non-linear proliferation response to progestins, we performed an RNA-Seq analysis of synchronized T47D cells following treatment with vehicle, low dose (0.1nM), or high dose (10nM) R5020 for 6hr or 18hrs. Evaluating the top 1,000 differentially expressed genes (p-adj<0.05) using K-means analysis, we identified three functionally distinct clusters of progestin-responsive genes (**Figure 2A**). Cluster 1 contains mRNAs whose progestin-dependent expression is regulated in a biphasic manner (e.g. *CCND1, E2F1, SGK1*), with maximal expression being induced in response to low dose (0.1nM) treatment as opposed to high dose (10nM) treatment (see expanded validation in **Figure S2A**). Cluster 2 contains mRNAs that exhibit a classical dose-dependent linear response to progestin treatment (e.g. *HSD17B2, S100A1, KLF4*). Cluster 3 contains mRNAs whose transcription is repressed by high dose progestin treatment (e.g. *FOXA1, ID1, CYP1A1*). The expression patterns of several genes in each cluster were validated using RT-qPCR in response to R5020 (**Figure 2B**) or to progesterone (P4) (**Figure S2B**).

**Figure 2.**
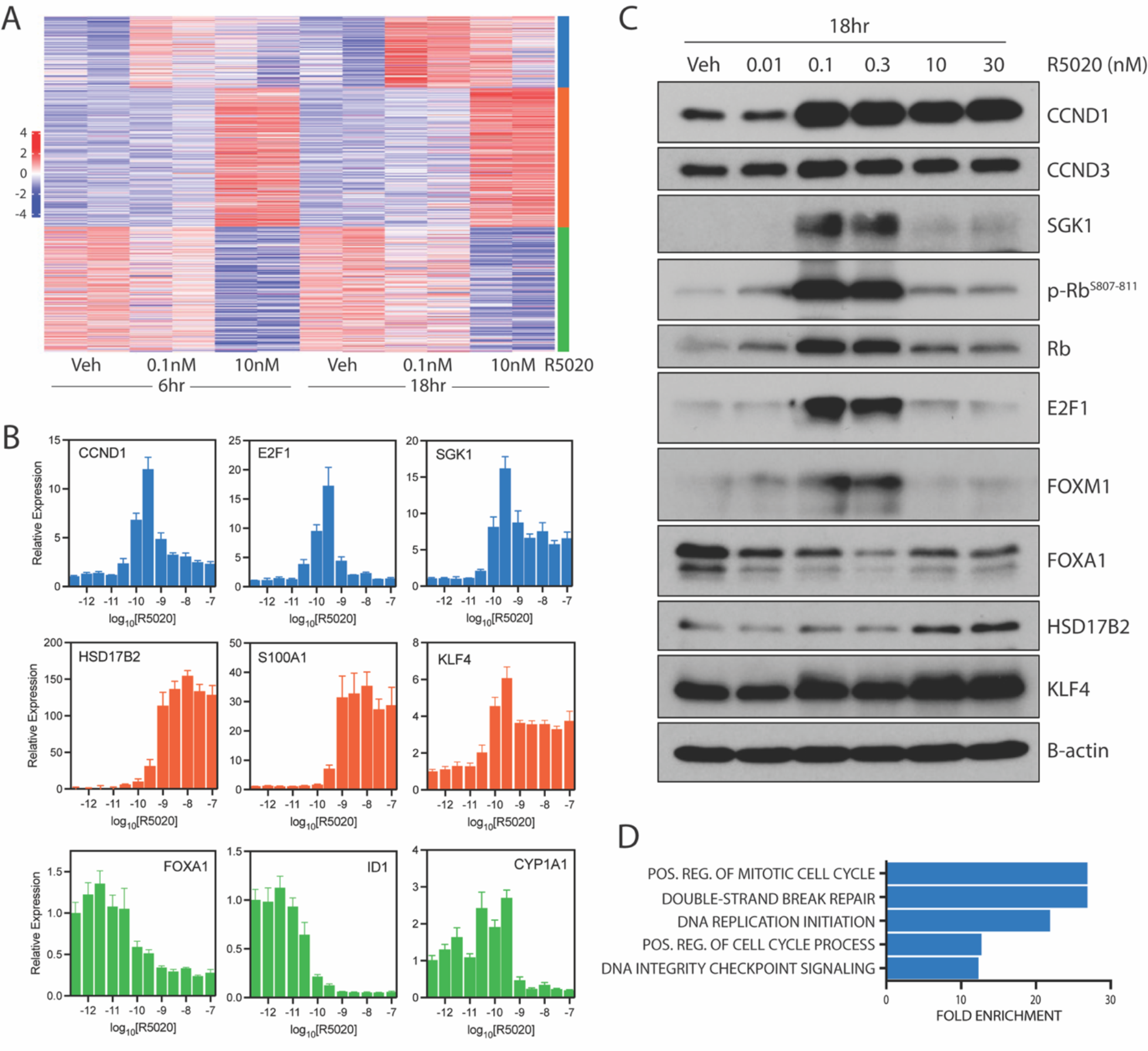
Low dose progestin treatment drives sustained cell cycle entry through expression of key cell cycle regulators. (A) Synchronized T47D cells were treated with vehicle, 0.1nM, or 10nM R5020 for 6hrs or 18hrs. Cells were harvested, and RNA was isolated for sequencing. K-means clustering analysis enabled the identification of three distinct clusters of DEGs. (B) Representative transcripts for each gene expression cluster were validated by RT-qPCR. Synchronized T47D cells were treated with R5020 for 18hrs, and their RNA was isolated for RT-qPCR analysis. Results are expressed as relative expression, calculated by the ≥≥Ct method ± standard devia+on. (C) Representative DEGs for each cluster were validated by SDS-PAGE with immunoblot. (D) Biphasic proges+n response genes are enriched for biological functions related to cell proliferation. DEGs identified in K-means cluster 1 (top) were subject to PANTHER Gene Ontology analysis. Top five significantly enriched biological function terms are shown.

The importance of these changes in transcription was confirmed by showing that the expression of cyclin D1 and E2F1 proteins mirrored that of their respective mRNAs, which was accompanied by a robust increase in phosphorylation of retinoblastoma (Rb) (**Figure 2C**). Follow-up analysis of the biphasic genes in cluster 1 using PANTHER Gene Ontology revealed that genes in this cluster were implicated in cell cycle regulation and DNA repair. The top five significantly enriched biological processes are shown in **Figure 2D**. Using the same strategy, we were unable to identify any significantly enriched biological processes in clusters 2 and 3. Thus, we conclude that low dose progestin treatment (0.1nM) alone drives the expression of key cell cycle regulators, which aligns with the observed phenotype of enhanced proliferation in T47D breast cancer cells. Consequently, we have determined that the proliferation of breast cancer cells in response to low doses of progestins can be attributed to specific effects on the expression of genes implicated in cell division.

### Low dose progestins activate the CCND1-Rb-E2F signaling axis

We were interested in identifying the primary regulators of the proliferative responses to progestins and therefore performed Ame course dose response studies to establish the temporal relationships of the expression of the genes implicated as mediators of the proliferative responses. To this end, synchronized T47D cells were treated with low (0.1nM) or high (10nM) dose R5020 for 1 to 24hrs. Using RT-qPCR, we evaluated the relative transcript abundance over Ame for several key cell cycle regulators (Cluster 1). In response to low dose R5020, we found that maximal *CCND1* and *SGK1* expression was apparent ∼6-8hr post-treatment, while maximal *E2F1* and *FOXM1* expression occurred at later Ame points (18-24hrs) (**Figure 3A**). Importantly, the earlier expression of SGK1 and CCND1 and later expression of E2F1 and FOXM1 was confirmed at the protein level using immunoblot analysis (**Figure 3B**). Enhanced CCND1 and SGK1 protein expression is detectable as early as 3hr post-treatment, in temporal alignment with the phosphorylation of Rb. Enhanced E2F1 and FOXM1 expression occurs later at 12 to 18hrs. Low-dose progestin-induced *CCND1* and *SGK1* mRNA expression is not sensitive to the translational inhibitor cycloheximide (CHX), indicating that these transcriptional responses are primary responses to progestin treatment (**Figure 3C**). In contrast, CHX blocks expression of *E2F1* and *FOXM1* mRNAs, indicating that these genes are secondary targets of the progestin response.

**Figure 3.**
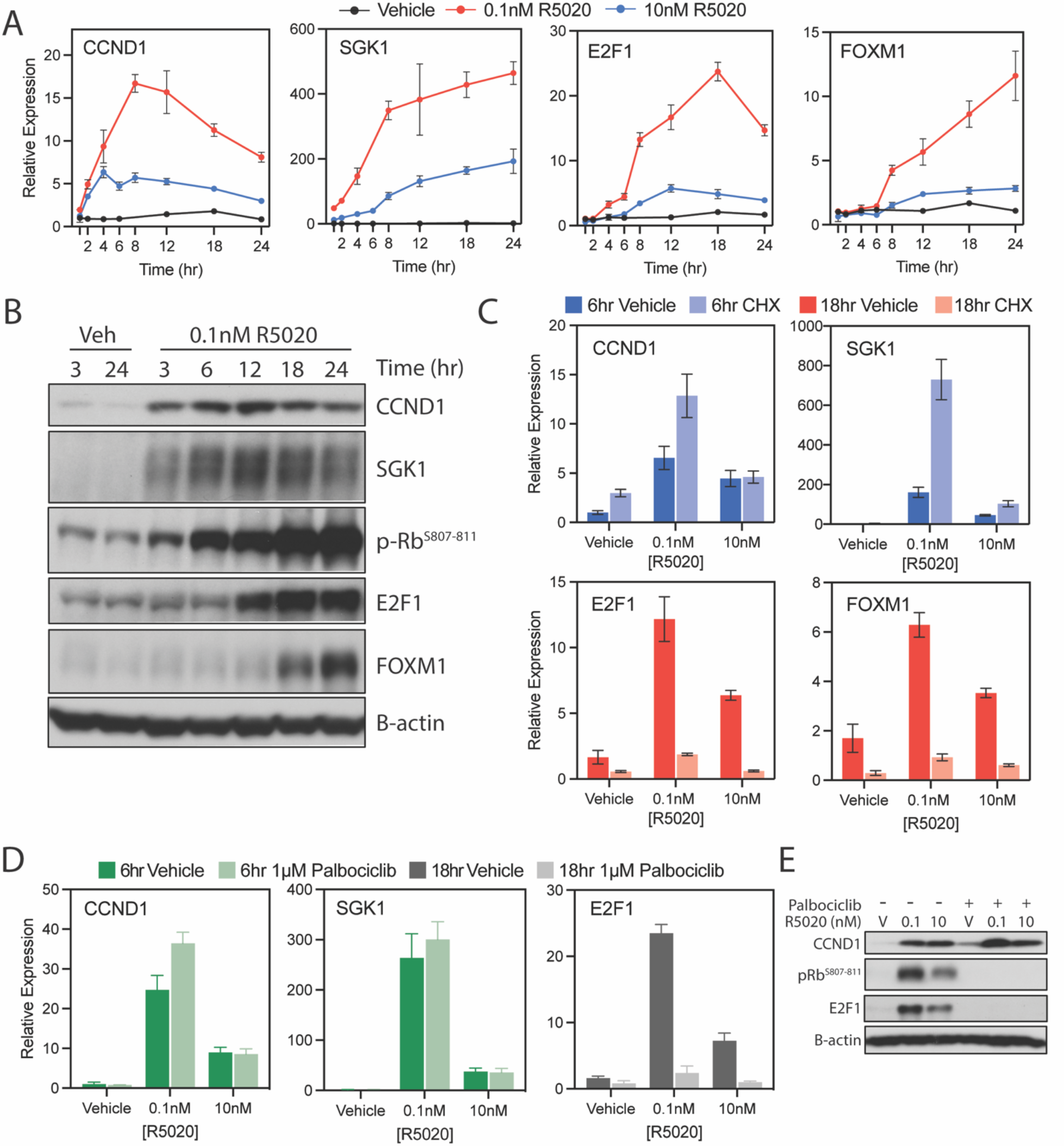
Low dose progestin treatment activates the CCND1-Rb-E2F signaling axis. (A) *CCND1* and *SGK1* mRNA expression temporally precedes that of *E2F1* and *FOXM1*. Synchronized T47D cells were treated with vehicle, 0.1nM R5020, or 10nM R5020. Following treatment for a range of times, RNA was isolated for RT-qPCR. Results are expressed as relative expression, calculated by the ΔΔCt method ± standard deviation. (B) CCND1 and SGK1 protein abundance are enhanced as early as 3hr, while E2F1 and FOXM1 protein expression arises later. Relative protein abundance was assessed over +me using SDS-PAGE with immunoblot. (C) *CCND1* and *SGK1* are insensitive to translational inhibition by cycloheximide, while *E2F1* and *FOXM1* expression is blocked. Synchronized T47D cells were treated with vehicle, 0.1nM R5020, or 10nM R5020 with or without 5ug/ml cycloheximide. RNA was isolated for RT-qPCR, and results are expressed as relative expression, calculated by the ΔCt method ± standard deviation. (D) *CCND1* and *SGK1* are insensitive to CDK4/6 inhibition by palbociclib, while *E2F1* and *FOXM1* expression is blocked. Synchronized T47D cells were treated with vehicle, 0.1nM R5020, or 10nM R5020 with or without 1μM palbociclib. RNA was isolated for RT-qPCR, and results are expressed as relative expression, calculated by the ΔCt method ± standard deviation. (E) CCND1 protein expression is insensitive to 1μM Palbociclib, while phosphoryla+on of Rb and E2F1 expression are blocked. Relative protein abundance was assessed over +me using SDS-PAGE with immunoblot.

It is well documented that the expression of cell cycle genes in breast cancer cells are facilitated by CCND1-cyclin-dependent kinase 4 and 6 (CDK4/6) dependent Rb phosphorylation, release of E2F proteins, and increased expression of E2F1 responsive genes (e.g. *FOXM1, CDC6*) (32). Accordingly, inhibition of CDK4/6 activity using palbociclib did not inhibit the expression of either CCND1 or SGK1 (mRNA or protein) (**Figure 3D-E**). In contrast, phosphorylation of Rb and the expression of *E2F1, CDC6*, and *FOXM1* do require CDK4/6 activity, as both activities were inhibited by palbociclib (**Figure S3C**). These results highlight the temporal importance of the CCND1-Rb-E2F in mediating proliferative responses to progestins but also introduce SGK1 as an early target of low dose PR action.

### Transcriptional response to low dose progestin treatment does not require canonical PR action

The observation that maximal transcriptional activity of *CCND1, SGK1*, and *E2F1* (and phosphorylation of Rb) occur at 0.1nM (R5020 or P4) was surprising given that at this concentration of ligand, the fractional occupancy of PR is exceptionally low. We have confirmed that PR is required for the responses observed using targeted *PGR* siRNA knock-down (achieving >80% knock-down – **Figure S4A-B**). Notably, targeted depletion of PR expression resulted in a near complete inhibition of progestin-mediated expression of *CCND1, SGK1*, and *E2F1* (**Figure 4A**). A genome wide ChIP sequencing study was conducted with the goal of identifying enhancers which were directly regulated by PR in response to low dose progestins. Surprisingly however, at doses of progestin which resulted in maximal induction of the expression of *CCND1* and *SGK1* mRNAs, PR chromatin binding events were exceptionally rare (at PREs or otherwise) (data not shown). The most contemporary models of PR action hold that the transcriptional activity of the receptor requires specific post-translational modifications. Specifically, MAPK-induced phosphorylation at Ser294 has been shown to be required for nuclear translocation and transcriptional activity on PRE-containing enhancers (33). Thus, we assessed phosphorylation of PR^S294^ in response to low and high dose R5020. However, we were unable to detect phosphorylation of PR at Ser294 at the levels of progestin required for maximal expression of *SGK1* and *CCND1* mRNAs (0.1-0.3nM) (**Figure 4B**). Thus, whereas PR is required for the expression of *CCND1* and *SGK1* mRNAs, the low levels of hormone required suggest that PR may be used in a non-canonical manner (e.g. DNA tethering, cytoplasmic signal transduction), as opposed to operating in a classical manner (PR/PRE dependent).

**Figure 4.**
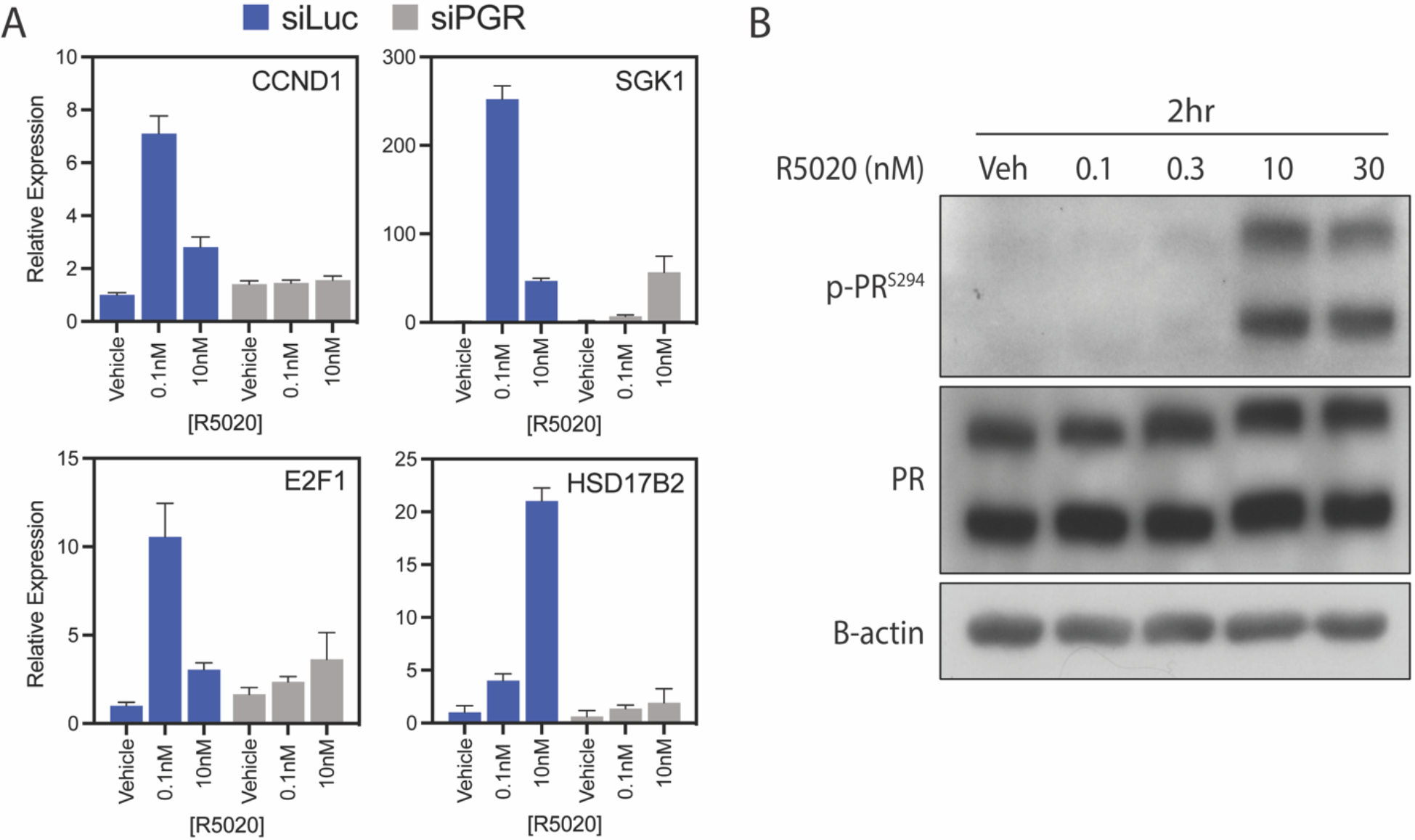
Classical PR action is not detected in response to low dose progestin treatment. (A) siRNA knockdown of *PGR* transcripts blocks progestin-mediated transcription of key target genes. T47D cells were reverse transfected with siRNA targeted to luciferase or *PGR*, followed by synchronization by serum deprivation. Synchronized transfected T47D cells were then treated with vehicle, 0.1nM R5020, or 10nM R5020 for 18hrs. RNA was isolated for RT-qPCR, and results are expressed as relative expression, calculated by the ΔCt method ± standard deviation. (B) Low dose R5020 does not result in increased PR phosphorylation at Ser294, a key marker of canonical PR activity. Synchronized T47D cells were treated with a range of R5020 doses for 2hrs. Cell lysate was then subjected to SDS-PAGE and immunoblot analysis.

### Cyclin D1 and SGK1 are required for low dose progestin-mediated CCND1-Rb-E2F axis activation

Our studies identify *CCND1* and *SGK1* as early, primary targets of PR in cells treated with low doses of progestins, and thus, we next evaluated the relative importance of these proteins on proliferative responses to this hormone. Using siRNA knock-down, we determined that *CCND1* expression is required for the expression of *E2F1*, but not for *SGK1* (**Figure 5A**). Further, as expected, the expression of the biphasic PR target gene *IGF1* is unaffected by *CCND1* knock-down. Knock-down of *SGK1* impairs low dose progestin-mediated *E2F1* expression, and unexpectedly was also shown to decrease the expression of *CCND1* (**Figure 5B**). *SGK1* knock-down does not attenuate biphasic response gene *IGF1* mRNA expression. Together, these data suggest that low dose progestin-mediated expression of *CCND1* and *SGK1* are somewhat codependent and that both are required for the optimal expression of the key cell cycle gene *E2F1*.

**Figure 5.**
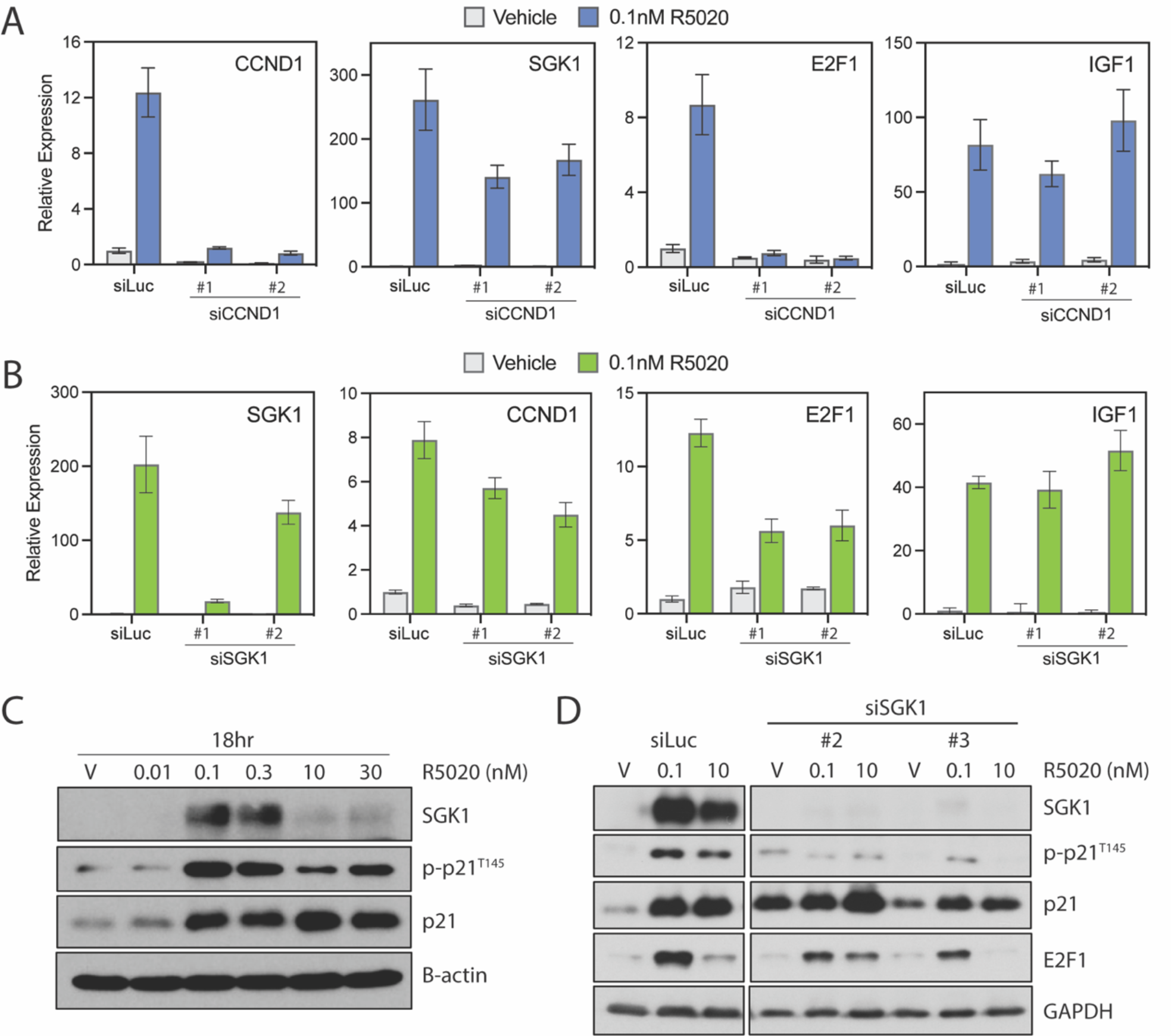
CCND1 and SGK1 are required for low dose progestin-mediated ac+va+on of the Rb-E2F axis. (A) siRNA knockdown of *CCND1* impairs progestin-mediated expression of *E2F1*. T47D cells were reverse transfected with siRNA targeted to luciferase or *CCND1*, followed by synchronization by serum deprivation. Synchronized transfected T47D cells were then treated with vehicle, 0.1nM R5020, or 10nM R5020 for 6hrs. RNA was isolated for RT-qPCR, and results are expressed as relative expression, calculated by the ΔΔCt method ± standard devia+on. (B) siRNA knockdown of *SGK1* impairs progestin-mediated expression of *E2F1*. T47D cells were reverse transfected with siRNA targeted to luciferase or *SGK1*, followed by synchronization by serum deprivation. Synchronized transfected T47D cells were then treated with vehicle, 0.1nM R5020, or 10nM R5020 for 8hrs. RNA was isolated for RT-qPCR, and results are expressed as relative expression, calculated by the ΔΔCt method ± standard devia+on. (C) SGK1 expression correlates closely with phosphorylation of p21 at Thr145. Synchronized T47D cells were treated with a range of R5020 doses for 18hrs. Cell lysate was then subjected to SDS-PAGE and immunoblot analysis. (D) siRNA knockdown of *SGK1* blocks progestin-mediated phosphorylation of p21 at Thr145 and partially acenuates E2F1 expression. T47D cells were reverse transfected with siRNA targeted to luciferase or *SGK1*, followed by synchronization by serum deprivation. Synchronized transfected T47D cells were then treated with vehicle, 0.1nM R5020, or 10nM R5020 for 18hrs. Cell lysate was then subjected to SDS-PAGE and immunoblot analysis.

Phosphorylation of the cyclin-dependent kinase inhibitor protein p21 (CDKN1A) impedes its anti-proliferative activity by excluding it from the nucleus (34). Although not previously described as a target of SGK1, we observed that this enzyme shares a serine/threonine kinase domain that is similar to that of AKT – a kinase known to phosphorylate p21 (34-36). Interestingly, we determined that while p21 protein expression in cells is increased by progestin (0.1-30nM), its phosphorylation at Thr145 (inhibitory) is robustly increased at even the lowest doses of hormone (0.1-0.3nM) when maximal expression of SGK1 is apparent (**Figure 5C**). Furthermore, targeted knock-down of *SGK1* by siRNA severely attenuates progestin-mediated phosphorylation of p21 at Thr145, absent a change in absolute p21 levels. We conclude that p21 is a phosphorylation target of SGK1 (**Figure 5D**). This is significant, as this particular phosphorylation event results in the exclusion of p21 from the nucleus, an activity that would disable its inhibitory actions on proliferation. Taken together, these results suggest that CCND1 and SGK1 act cooperatively to enable activation of CCND1-CDK4/6 activation while disabling the anti-proliferative actions of p21.

## DISCUSSION

Progesterone levels fluctuate dramatically during development and during the normal reproductive cycle in females, and it is well established that there are non-overlapping phenotypic responses to different levels of this hormone. It is thus implied that cells possess the ability to recognize and respond to different levels of hormone, although the mechanisms that enable this non-linear pharmacology have not been defined. What has been inferred is that low and high dose biology reflects quantitative, dose proportional differences in the expression of a common set of genes. The results of the studies herein would argue against such a model in favor of one in which posits that progestins exhibit a non-linear pharmacology with low and high dose progestins driving the expression of very different transcriptomes, resulting in different phenotypic responses. Of particular importance is the observation, in an established cellular model of breast cancer, that low dose progestins induce the expression of mRNAs whose encoded proteins are required for cell proliferation. Interestingly, most of these genes are likely to be secondary responses to progestins that result from primary regulatory events which enable the phosphorylation of Rb (and the subsequent activation of E2F1 signaling) and inactivation by phosphorylation of the cell cycle inhibitor p21. As we and others have reported in the past, the transcriptional upregulation of *CCND1* expression by PR/low dose progestins results in increased CDK4/6 activity and hyperphosphorylation of Rb. What was unexpected was our finding that the expression of SGK1, also a primary response to low dose progestins, results in increased phosphorylation of p21 – a key negative regulator of the cell cycle. Inhibition of either SGK1 or CCND1 expression is su?cient to inhibit progestin-mediated expression of *E2F1*, and importantly we find that both genes are regulated by low dose progestins (**Figure 6**). What remains to be determined is how, considering the classical models of PR pharmacology, the induction of *SGK1* and *CCND1* expression can be accomplished by doses of progestins which allow only a fractional occupancy of PR in the cell. Previously, it has been demonstrated that phosphorylation of PR-B at Ser294 reads on the nuclear translocation and transcriptional competency of this receptor (33). However, at the doses of progestins required for proliferation, and for maximal induction of CCND1/SGK1 expression, we were unable to detect phosphorylated PR Ser294. These data suggest either that the technologies we have available to study PR-enhancer interactions and PR phosphorylation are not sensitive enough to detect the small fraction of nuclear PR that drives these proliferative responses and/or that non-canonical actions of the receptor in the cytoplasm contribute to these primary responses to low dose progestins. Recently it has been demonstrated that extremely low a?nity estrogens, which achieve only transient estrogen receptor (ER) occupancy enable the quantitative activation of MAPK and mTOR signaling in the cytoplasm (37). Thus, it is possible that similarly low doses of progestins may regulate activities in the cytoplasm that contribute to its proliferative actions.

**Figure 6.**
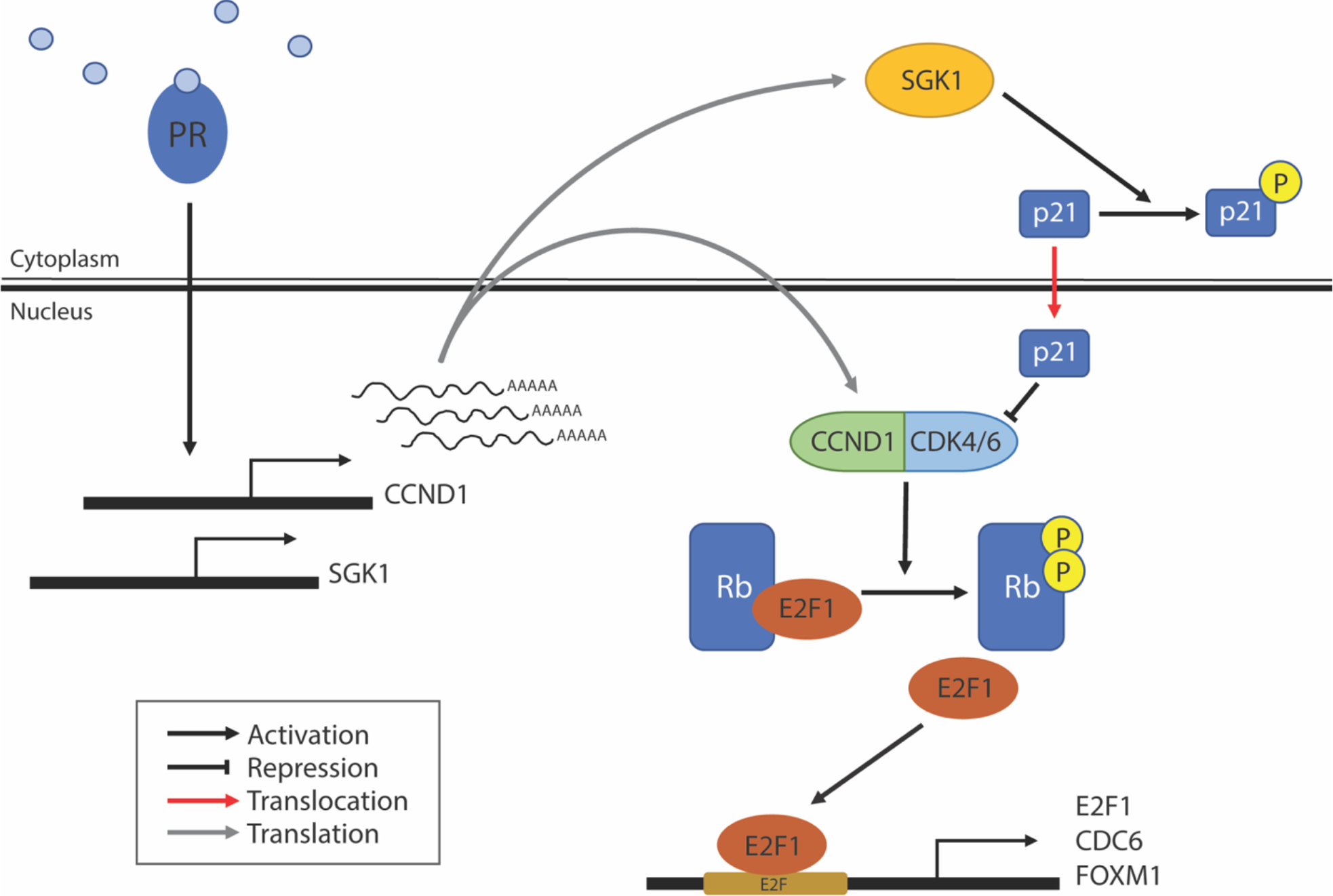
Proposed molecular mechanism for low dose progestin-mediated proliferation in T47D breast cancer cells. PR drives enhanced expression of *CCND1* and *SGK1*, which are translated into protein to perform complementary functions in activating the Rb-E2F signaling axis. CCND1 cooperates with CDK4/6 to phosphorylate Rb, thereby releasing E2F1 to act on key downstream promoter regions. In complement, SGK1 phosphorylates p21, thereby excluding it from the nucleus and blocking its inhibitory function on CDK4/6.

The primary focus of this study was to define the mechanisms underlying the low dose biology of progestins. However, we observed that at the doses of progestins required to induce the expression of the classical PR-target genes (e.g. *S100A1, HSD17B2*), the low dose responsive genes (and proliferation) are generally repressed. We have not yet explored the mechanisms by which high doses of progestins inhibit proliferation/proliferative responses, although our data are compatible with the idea that dimeric nuclear PR is involved in the acAve repression of low dose responsive genes. It is notable that we observe downregulation of FOXA1, a protein required for nuclear receptor action at most enhancers, in cells treated with high dose progestins. Further exploration of the mechanisms by which high dose progestins regulate gene repression will likely be informative as to the mechanisms underlying the e?cacy of progestins when used as therapies in advanced ER-positive breast cancer.

The Women’s Health Initiative (WHI) was a very large clinical study that had the primary objective of assessing the effects of supplemental hormone therapy on cardiovascular health, osteoporosis and the risk of breast and colon cancer (38). The study was neutral with respect to cardiovascular health and positive with respect to osteoporosis. However, a small but significant increase in breast cancer risk was observed in women who took conjugated equine estrogens (CEE) together with a progestin (MPA) (PREMPRO), as indicated in women with an intact uterus. A sub study was performed in hysterectomized women who could take estrogens without a progestin, and it was observed that breast cancer risk (and colon cancer risk) was dramatically reduced (39). These and other studies have led to the conclusion that it was the progestin that was responsible for the negative impact on the breast. Whereas it has been assumed that it was the androgenic activity of MPA that contributed to breast cancer pathobiology, the results of our studies suggest that it may also relate to the dose of the progestin used. The dose of MPA in PREMPRO that was evaluated in women with an intact uterus was determined based on the minimum amount of the progestin required to protect the uterus. It is therefore possible that some women were exposed to doses of the progestin that were proliferative (i.e. low doses). Indeed, increased breast density, an established risk factor for breast cancer, is increased in some patients on the standard dose PREMPRO (40). There are ongoing studies exploring the breast cancer risk in patients taking 17β-estradiol with natural P4 (no androgenic activity). The results of these studies may be instructive as to the best and safest way to protect the uterus in women taking estrogen containing HRT.

Progestins remain an important component of contraceptives and hormone-based therapies to treat the climacteric patient. The studies we have performed, albeit confined to a model of breast cancer, have highlighted an unexpected complexity in PR pharmacology that should be considered in the delivery of existing medicines and in the development of new agents that target PR in cancer and in other endocrinopathies.

## METHODS

### Tissue culture

Immortalized human breast carcinoma T47D cells were maintained in RPMI 1640 supplemented with 8% fetal bovine serum (FBS), MEM Non-Essential Amino Acids (NEAA), and Sodium Pyruvate (NaPyr). Cell plates were incubated at 37°C in the presence of 5% CO_2_, and cultures were passaged 1-2 Ames weekly. Cells were routinely screened for mycoplasma contamination via PCR test.

### Proliferation assay

T47D cells were plated 2x10^6^ cells per 10cm culture dish for cell cycle synchronization in preparation for proliferation assays. Cells were allowed to attach overnight in phenol red-free RPMI 1640 supplemented with 8% charcoal stripped fetal bovine serum (CFS), NEAA, and NaPyr. Synchronization was achieved by 24hr serum starvation in phenol red-free RPMI 1640 supplemented with 0.1% CFS, NEAA, and NaPyr. Synchronized T47D cells were plated 10,000 cells per well on 48-well culture plates (avoiding the edge wells for evaporation risk) in RPMI 1640 supplemented with 8% CFS or 8% FBS, NEAA, and NaPyr. Cells were allowed to attach for 1-2hr, at which Ame they were treated with the relevant assay compounds. Due to compound instability in culture conditions, plates treated with P4 received 50% volume media replacements every 48hr to maintain proper drug concentration. Cell plates were incubated at 37°C in the presence of 5% CO_2_ for 7 days or until wells approach confluence, whichever is reached first. Upon assay completion, media was removed by plate inversion, and plates were frozen at -80°C overnight. Frozen plates were thawed to room temperature, and 200ul of ultrapure water was added per well. Plates were incubated for 2-3hr at 37°C prior to a second overnight freeze at -80°C. Plates were again thawed to room temperature and 200ul TNE buffer (NaCl, Tris, EDTA) + Hoechst dye (1.0 mg/ml, 1:500) was added to each well. DNA content, a surrogate for cell number, was measured using a plate reader by fluorescent absorbance at 460nm. Each treatment condition was assessed in triplicate, and results are expressed as relative fluorescent units +/-SD.

### Cell cycle analysis

T47D cells were plated 2.5x10^5^ cells per well in 6-well culture plates. Cells were allowed to attach overnight in phenol red-free RPMI 1640 supplemented with 8% CFS, NEAA, and NaPyr. Synchronization was achieved by 48hr serum starvation in phenol red-free RPMI 1640 supplemented with 0.1% CFS, NEAA, and NaPyr. Synchronized T47D cells were then replaced in phenol red-free RPMI 1640 supplemented with 8% CFS, NEAA, and NaPyr and treated with the relevant assay compounds. Cell plates were incubated at 37°C in the presence of 5% CO_2_ for 18hr. Upon treatment completion, media was removed by aspiration, and cells were gently washed with sterile PBS. Intact cells were gently dissociated from the culture plate using 0.05% trypsin, which was followed by washing with PBS. Cells suspended in PBS were then fixed by ice cold 100% ethanol, added dropwise to a final concentration of 70%. Fixed cells were stored at -20°C overnight. The 70% ethanol was removed, and cells were resuspended in cold PBS containing 50ug/ml propidium iodide and 100ug/ml RNase. Samples were assessed by flow cytometry within 48hr. Using forward and side scatter data, cell debris was gated out and single cells were assessed for their fluorescence intensity. Gates defining G1, S, and G2 phases were assigned using vehicle treated cells.

### RNA isolation and real-time quantitative PCR

T47D cells were plated 2.5x10^5^ cells per well in 6-well culture plates. Cells were allowed to attach overnight in phenol red-free RPMI 1640 supplemented with 8% CFS, NEAA, and NaPyr. Synchronization was achieved by 24-48hr serum starvation in phenol red-free RPMI 1640 supplemented with 0.1% CFS, NEAA, and NaPyr. Synchronized T47D cells were treated with the relevant assay compounds. Cell plates were incubated at 37°C in the presence of 5% CO_2_ for the noted amount of Ame. Upon treatment completion, media was removed by aspiration, and cells were gently washed with sterile PBS. Isolation of total RNA was performed using the Aurum™ Total RNA Mini-Kit according to the manufacturer’s instructions (Bio-Rad, Hercules, CA). Total RNA concentration and quality were determined using a Nanodrop 1000 (ThermoFisher). The iScript™ cDNA synthesis Kit (Bio-Rad) was used to reverse transcribe total RNA (1ug). Resulting cDNA was diluted 1:20 for further use in quantitative PCR analysis. qPCR was performed using 2ul of Bio-Rad SYBR green master mix with 65nM forward and reverse primer mix plus 1.26ul of diluted cDNA, resulting in a total reaction volume of 3.26ul. Each sample was assessed in triplicate. PCR amplification was performed using the Bio-Rad CFX384 qPCR system. Expression of *36B4* was used as a normalization control, and fold change expression for each target transcript was calculated as 2^-ΔΔCt^. Transcript-specific primers were purchased from Eton Biosciences, Inc.

### RNA Sequencing (RNA-Seq)

Read alignment, quality control and data analysis for duplicate samples were performed using Visualization Pipeline for RNA-seq (VIPER) (41). Alignment to the hg19 human genome was done using STAR v2.7.0f (42) followed by transcript assembly using cu?inks v2.2.1 (43) and RseQC v2.6.2 (44). Differential gene expression analyses were performed on absolute gene counts for RNA-Seq data and raw read counts for transcriptomic profiling data using DESeq2v1.18.1 (45). Hierarchical k-means clustering was performed using normalized gene expression representing the 1000 most variable genes between treatment samples versus control samples. Hierarchical k-means clustering was implemented by the kmeans function in the stats R package with k=3. Heatmap visualization was done by R package ComplexHeatmap (46).

### Immunoblot analysis

T47D cells were plated 2x10^6^ cells per 10cm culture dish. Cells were allowed to attach overnight in phenol red-free RPMI 1640 supplemented with 8% charcoal stripped fetal bovine serum (CFS), NEAA, and NaPyr. Synchronization was achieved by 24-48hr serum starvation in phenol red-free RPMI 1640 supplemented with 0.1% CFS, NEAA, and NaPyr. Whole-cell extracts were prepared using RIPA buffer supplemented with protease inhibitors and phosphatase inhibitor (50 mM Tris pH 7.5, 200 mM NaCl, 1.5mM MgCl2, 1% Triton X-100, 1mM EGTA, 10% glycerol, 50mM NaF, 1X protease inhibitor cocktail, 2mM sodium orthovanadate). Concentration of cleared cell lysate was assessed by the Pierce BCA Protein Assay Kit (ThermoFisher). Equal amounts of protein (20-40ug) were resolved via SDS-PAGE (8-10% polyacrylamide gels), followed by transfer to PVDF membrane and immunoblotting. The following antibodies were used at 1:1000 dilution: CCND1 (Invitrogen #MA5-14512), CCND3 (Cell Signaling #2936), SGK1 (Cell Signaling #121035), phosoho-Rb^Ser807-811^ (Cell Signaling #8516), Rb (Cell Signaling #9309), FOXM1 (Cell Signaling #204595), FOXA1 (Abcam #ab55178), HSD17B2 (Proteintech #10978-1-AP), KLF4 (Cell Signaling #4038), PR (Invitrogen #MA1-410), phospho-PR^Ser294^ (Cell Signaling #13736), p21 (Santa Cruz Biotechnologies #sc-397), phospho-p21^Thr145^ (Invitrogen #PA5-12646), and ?-acAn (Sigma-Aldrich #A5441). AnA-E2F1 antibody (BD Pharmingen #554213) was used at 1:500 dilution.

