## Supplemental Figures for "Breast cancer cells can recognize and respond to different levels of progestins to achieve different phenotypic outputs"

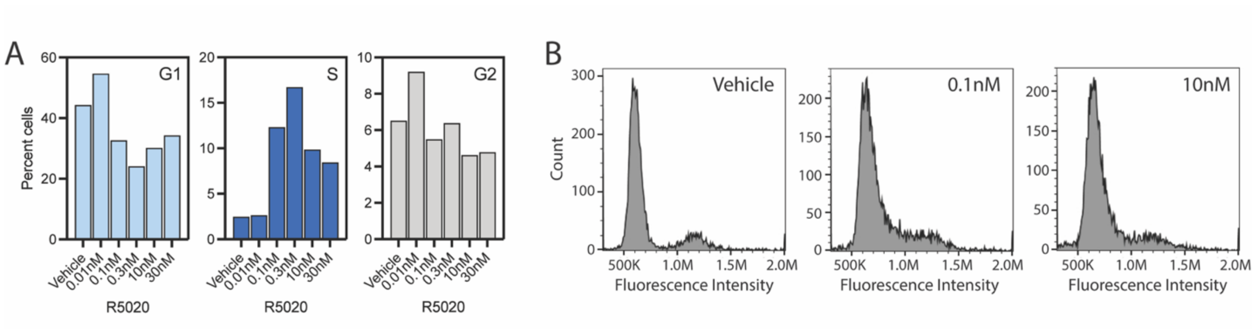


**Supplemental Figure 1.** Low dose progestins drive cell cycle entry. Synchronized T47D cells, grown in CFS, were treated with different doses of R5020 for 18hr. Cells were fixed and stained with DNA stain propidium iodide and assessed by flow cytometry. (A) Percent cell population in G1, S, and G2 phases of the cell cycle following 18hr R5020 treatment, as measured by propidium iodide. (B) Representative histograms depicting cell number and DNA content as measured by propidium iodide flow cytometry following 18hr treatment with R5020.

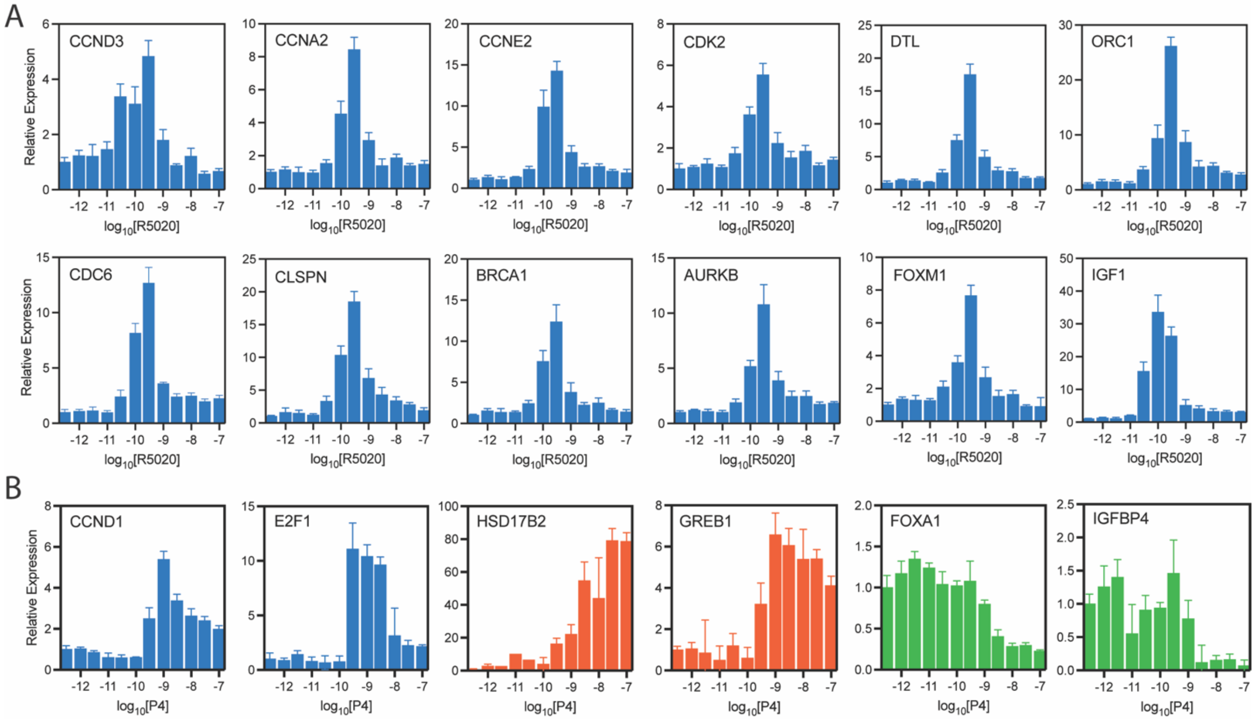


**Supplemental Figure 2.** (A) Expanded RT-qPCR validation of biphasic progestin dose response genes in the biphasic K-means cluster 1. Results are expressed as relative expression, calculated by the $\Delta\Delta$Ct method $\pm$ standard deviation. (B) Representative transcripts for each gene expression cluster were validated by RT-qPCR in response to P4. Synchronized T47D cells were treated with P4 for 18hrs, and their RNA was isolated for RT-qPCR analysis. Results are expressed as relative expression, calculated by the $\Delta\Delta$Ct method $\pm$ standard deviation.

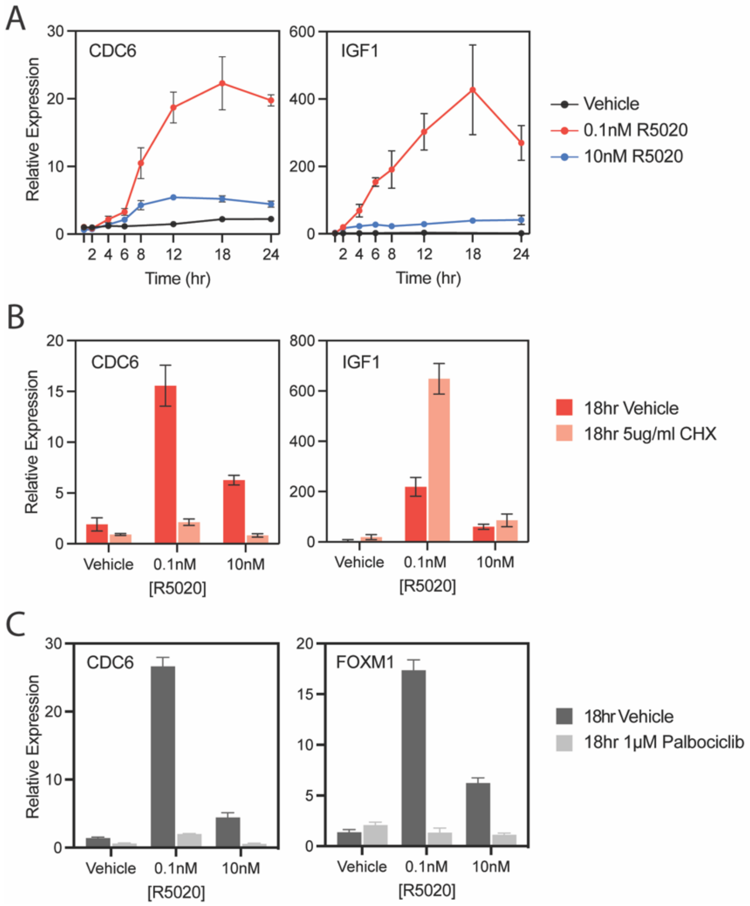


**Supplemental Figure 3.** (A) Expanded Figure 3A. (B) Expanded Figure 3C. (C) Expanded Figure 3D.

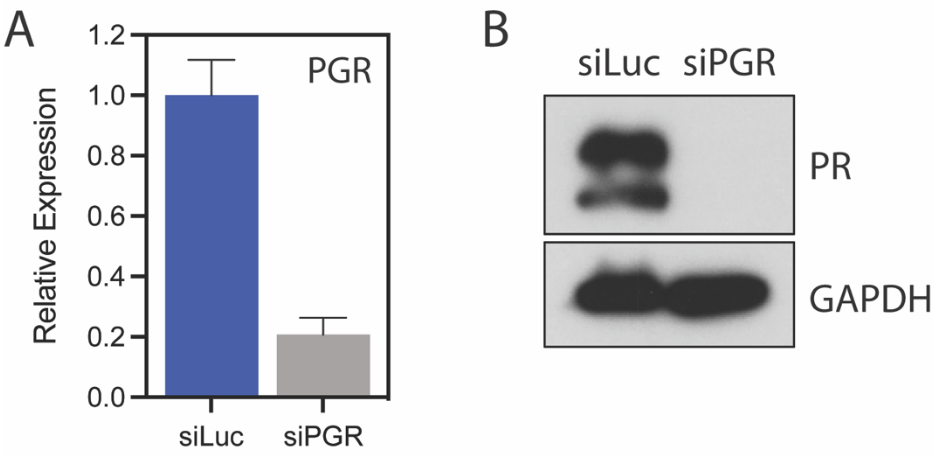


**Supplemental Figure 4.** Targeted siRNA of *PGR* resulted in >80% knock-down efficiency. Knock-down was validated by RT-qPCR (A) and immunoblot (B).
